# Riluzole as a Dual-Targeted Radiosensitizer for Osteosarcoma: Targeting Tumor Cells and Angiogenic Vasculature to Enhance Single High Dose Radiotherapy Efficacy

**DOI:** 10.1101/2025.10.07.681036

**Authors:** Pooja Prakash Rao, Charis Herbert, Syeda Maryam Azeem, Elena Gary, Gloria Ho, Raisa Munira, Hadi Askarifirouzja, Adriana Haimovitz-Friedman, Shahana Sultana Mahajan

## Abstract

Osteosarcoma is a highly aggressive bone malignancy primarily affecting children and young adults. It presents significant treatment challenges due to its inherent resistance to conventional fractionated radiotherapy (CFRT). Single high dose radiation therapy (SDRT) has promise for the treatment of radioresistant sarcomas, especially those characterized with extensive vascularity. However, its clinical application is severely constrained by toxicity to adjacent critical tissues. Radiosensitizers can enhance tumor cell susceptibility to radiation-induced DNA damage, improving therapeutic efficacy and potentially reducing collateral toxicity. Monotherapies targeting tumor vasculature alone in solid tumors have shown limited success as radiosensitizers in clinical settings. This highlights the importance of compounds that can simultaneously target both tumor cells and its associated microvasculature to maximize the therapeutic outcome to SDRT. Riluzole, the FDA-approved drug for Amyotrophic Lateral Sclerosis, is currently under investigation as a therapeutic agent for osteosarcoma. Riluzole acts to inhibit glutamate release, reduce glutathione levels in cancer cells, and mitigate tumor angiogenesis, positioning it as a potent radiosensitizing agent for the treatment of osteosarcoma. We hypothesize that Riluzole enhances osteosarcoma radiosensitivity to SDRT by simultaneously targeting intrinsic tumor radioresistance and pro-angiogenic signaling. Our findings demonstrate that Riluzole radiosensitizes osteosarcoma cells *in vitro* by reducing clonogenic survival and enhancing apoptosis. Mechanistically, Riluzole potentiates irradiation-induced reactive oxygen species (ROS) production, induces G2/M phase cell cycle arrest, inhibits DNA repair, and thereby amplifies radiation-induced DNA damage. Additionally, Riluzole suppresses radiation-induced Vascular Endothelial growth factor A (VEGFA) expression indicating its ability to overcome endothelial cell mediated radioresistance. Collectively, these results establish Riluzole as a promising radiosensitizer for osteosarcoma, with the potential to improve SDRT efficacy by overcoming both tumor-intrinsic and microvasculature-mediated radioresistance.

**Graphical abstract:** 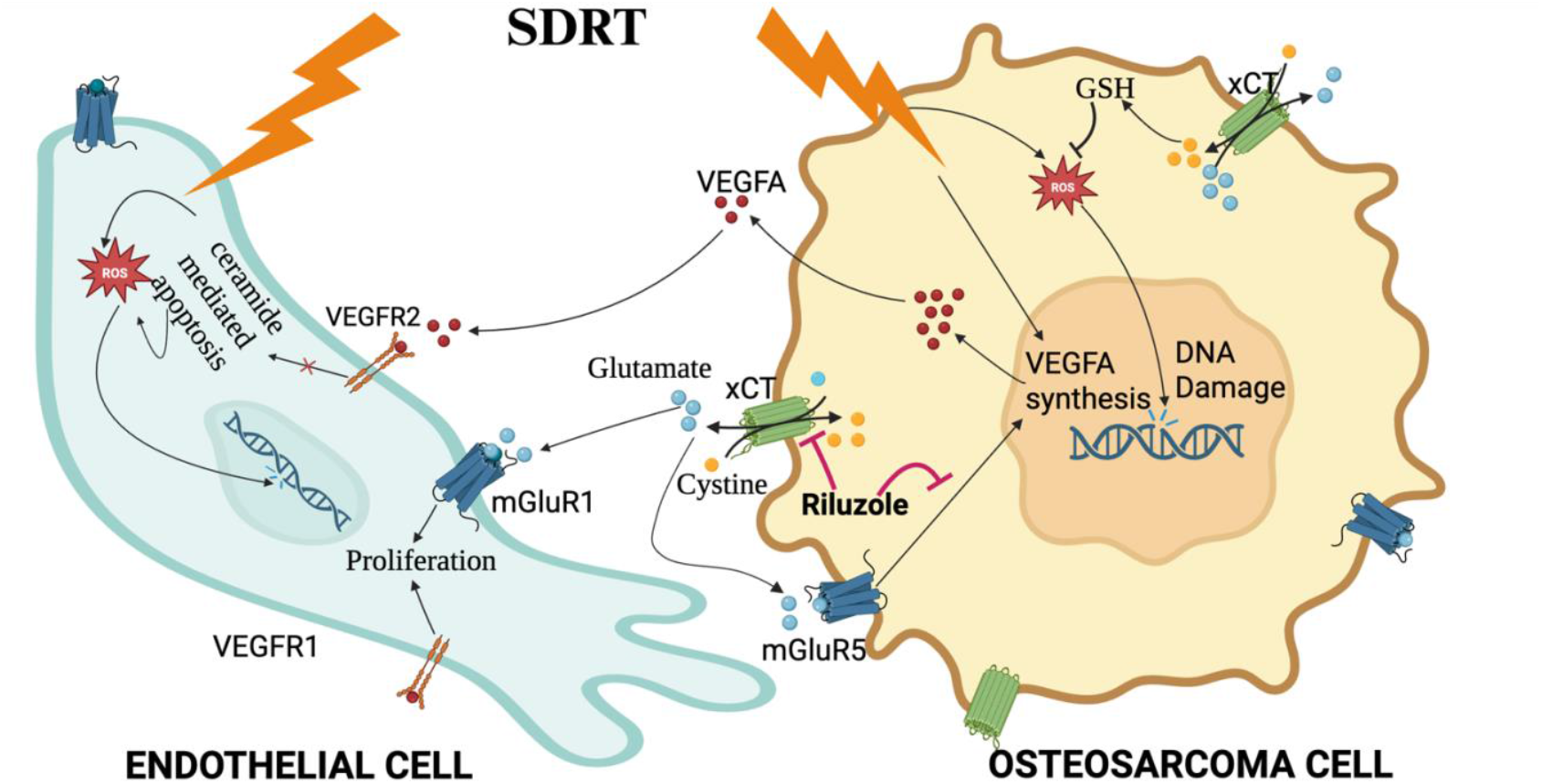

This schematic illustrates the proposed mechanism by which Riluzole enhances SDRT efficacy in osteosarcoma by targeting both tumor cells and VEGFA-mediated pro-survival signaling in endothelial cells. Riluzole increases radiation-induced ROS levels, induces G2/M cell cycle arrest, and inhibits DNA repair in osteosarcoma cells, thereby overcoming intrinsic tumor radioresistance. It also suppresses tumor cell VEGFA expression, which may contribute to reduced pro-survival signaling in the angiogenic endothelial cells within the tumor microenvironment. Together, these effects sensitize osteosarcoma tumors to SDRT, improving therapeutic outcomes (Illustration created using BioRender (BioRender.com, 2025)).

## Introduction

Osteosarcoma is a rare, aggressive and radioresistant cancer of bone mostly seen in children and young adults^1,2^. Even though surgery followed by neoadjuvant chemotherapy is the standard first line treatment for osteosarcoma, radiation therapy (RT) still represents an important treatment modality especially for unresectable tumors or when limb amputation surgery is refused by the patients^3,4,5^. Further, the conventional treatment approaches have not been completely successful in preventing the metastasis of the tumor in 30-40% of patients, leading to poor prognosis and a low 5-year survival rate for osteosarcoma. The mechanisms underlying the intrinsic radioresistance nature of osteosarcoma are not yet fully understood, although several factors have been implicated. These include an enhanced capacity to repair radiation-induced DNA damage, deregulation of cell cycle control, and the hypoxic and hyper-vascular tumor microenvironment, characterized by aberrant angiogenesis and vascular signaling^6–11^. The limited success of conventional fractionated radiotherapy (CFRT) in osteosarcoma highlights the need for alternative treatment approaches. Elucidating the mechanisms that underlie osteosarcoma radioresistance and exploiting vascular dependencies may provide a foundation for improving survival outcomes and establishing novel therapeutic modalities.

Radiation therapy mediates tumor control mainly through the induction of DNA damage. The DNA damage response determines the cell fate by coordinating DNA repair and regulating cell cycle arrest. Approximately two-thirds of radiation-induced damage is mediated by reactive oxygen species (ROS)^12,13^. While excessive ROS can lead to cell death, cancer cells maintain elevated ROS levels without lethal damage by upregulating antioxidant defenses such as glutathione (GSH)^14–16^. Therapeutic strategies that disrupt this balance, such as inhibiting GSH synthesis have been explored to promote ROS accumulation, amplify DNA damage, and enhance radiosensitivity^17^.

Osteosarcoma cells exhibit robust DNA damage repair capacity, with poly(ADP-ribose) polymerase 1 (PARP1) playing a central role in repairing single-strand breaks (SSBs) during S and G2 phases^18^. Unrepaired SSBs can collapse replication forks, generating double-strand breaks (DSBs) that require homologous recombination (HR). Agents that induce G2/M arrest can synergize with PARP1 inhibition to promote DSB accumulation, impair DNA repair, and trigger apoptosis, representing a potent radiosensitization strategy. Furthermore, mutational landscape of osteosarcomas resembles that of BRCA-deficient tumors, further supporting the rationale for combining radiation with DNA repair inhibitors such as PARP1 inhibitors to overcome radioresistance^9^.

Single High-Dose Radiation Therapy (SDRT) is a specialized type of stereotactic body radiation therapy (SBRT) that delivers a single, high-dose fraction (>8 Gy) to the tumor. This offers multiple advantages over CFRT, in several cancer types including radioresistant sarcomas^11,19,20^. This is attributed to its ability to induce apoptosis in both tumor and endothelial cells, which are otherwise not effectively targeted by CFRT^10,21,22^. In osteosarcoma, microvascular density critically influences tumor response to radiation. Vascular endothelial growth factor A (VEGFA) is the most potent pro-angiogenic molecule and is one among the 13 most highly expressed genes in osteosarcoma^23–25^. VEGFA is known to play a central role in mediating tumor resistance to RT^26,27^. Importantly, tumors may respond to SDRT by upregulating VEGFA expression in both cancer cells and angiogenic endothelial cells, a response that can enhance endothelial survival, promote angiogenesis, and contribute to radioresistance^26,28^. VEGFA secreted by tumor cells can indirectly inhibit acid sphingomyelinase (ASMase)– mediated ceramide generation in endothelial cells. This suppression prevents ceramide-induced apoptosis triggered by SDRT, thereby promoting endothelial cell–mediated tumor radioresistance and limiting the maximal therapeutic efficacy of SDRT. ^29^

The efficacy of SDRT is further constrained by tumor size, anatomical location, and toxicity to surrounding normal tissues. Radiosensitizers have the potential to enhance the therapeutic efficacy of SDRT by broadening the therapeutic window and increasing tumor eradication^30,31^. While anti-angiogenic agents have been explored to overcome endothelial cell–mediated radioresistance, their clinical benefit in solid tumors has remained modest— largely due to compensatory signaling and reciprocal crosstalk between tumor cells and the tumor vasculature within the microenvironment^32,33^. These adaptive survival pathways limit the durability of vascular targeting strategies alone. Consequently, therapeutic strategies that simultaneously target intrinsic tumor cell radioresistance and angiogenic endothelial survival hold promise to achieve durable tumor regression and prevent recurrence following SDRT. Notably, no clinically approved radiosensitizers currently exist for osteosarcoma, underscoring an urgent need to develop innovative, mechanism-based strategies to enhance SDRT efficacy in this radioresistant malignancy.

Riluzole (6-trifluoromethoxy-2-benzothiazolamine) is an FDA-approved drug for Amyotrophic Lateral Sclerosis (ALS) that has demonstrated anticancer activity in multiple malignancies, including melanoma, glioma, breast, and prostate cancers^34,35^. Although its precise anticancer mechanism remains under investigation, Riluzole is reported to suppress glutamate signaling by reducing glutamate release from cancer cells, thereby preventing activation of glutamate receptor mediated signaling that drive tumor progression^34^. It has also shown to exhibit anti-angiogenic potential, particularly in glutamate-dependent tumors like breast cancer and has been shown to inhibit VEGF-induced endothelial cell proliferation and abnormal vessel formation in pre-clinical models^36,37^. Riluzole in combination with chemotherapy or radiotherapy has been reported to potentiate tumor responses, improving therapeutic efficacy^34^. Riluzole has been shown to sensitize metastatic melanoma and nasopharyngeal carcinoma to ionizing radiation^38,39^.

Given the critical role of glutamate signaling in regulating redox homeostasis, DNA repair, and angiogenesis — key drivers of radioresistance, Riluzole represents a promising radiosensitizer for osteosarcoma when combined with SDRT. Combining Riluzole with SDRT has the potential to enhance tumor control by simultaneously targeting tumor cells and vasculature while reducing the effective therapeutic dose of radiation. We hypothesize that Riluzole enhances osteosarcoma radiosensitivity by suppressing tumor-intrinsic resistance mechanisms and by modulating pro-survival signaling in endothelial cells. Our results show that Riluzole augments radiation-induced ROS generation, induces G2/M arrest and impairs DNA repair, and, when combined with SDRT, significantly decreases clonogenic survival and promotes apoptosis in osteosarcoma cells. Furthermore, by suppressing VEGFA expression in osteosarcoma cells, Riluzole may indirectly inhibit pro-survival signaling in endothelial cells. Together, these effects suggest that combining Riluzole with SDRT could permit lower effective radiation doses, broaden the therapeutic window and improve therapeutic outcomes in osteosarcoma.

## Results

### A. Riluzole Enhances Tumor Cell Radiosensitivity

#### Riluzole Reduces Survival of Irradiated Osteosarcoma Cells

To determine the effect of Riluzole on survival fraction of irradiated osteosarcoma cells, LM7 and OS482 cells were pre-treated with 25 μM Riluzole or DMSO (control) for 24 h and subjected them to varying dose of X-ray radiation (2 Gy - 8 Gy). *In vitro* clonogenic assay was used to assess the ability of Riluzole to radiosensitize osteosarcoma cells. As shown in the dose–survival curves **(Fig. 1)**, Riluzole pretreatment significantly reduced the survival fraction of irradiated osteosarcoma cells compared with radiation alone. In LM7 cells, high-dose irradiation (6 Gy and 8 Gy) induced modest cell death, consistent with their intrinsically radioresistant phenotype; However, pretreatment with Riluzole markedly enhanced radiosensitivity, resulting in significantly lower survival fractions at these doses compared to radiation alone **(Fig. 1A)**. In OS482 cells, Riluzole treatment significantly decreased clonogenic survival compared to controls, reflecting their relatively higher sensitivity to the drug. Moreover, across all radiation doses tested (2 Gy–8 Gy), Riluzole pretreatment further reduced survival fractions compared to radiation alone, further demonstrating its radiosensitizing potential **(Fig.1B)**.

**Figure 1:**
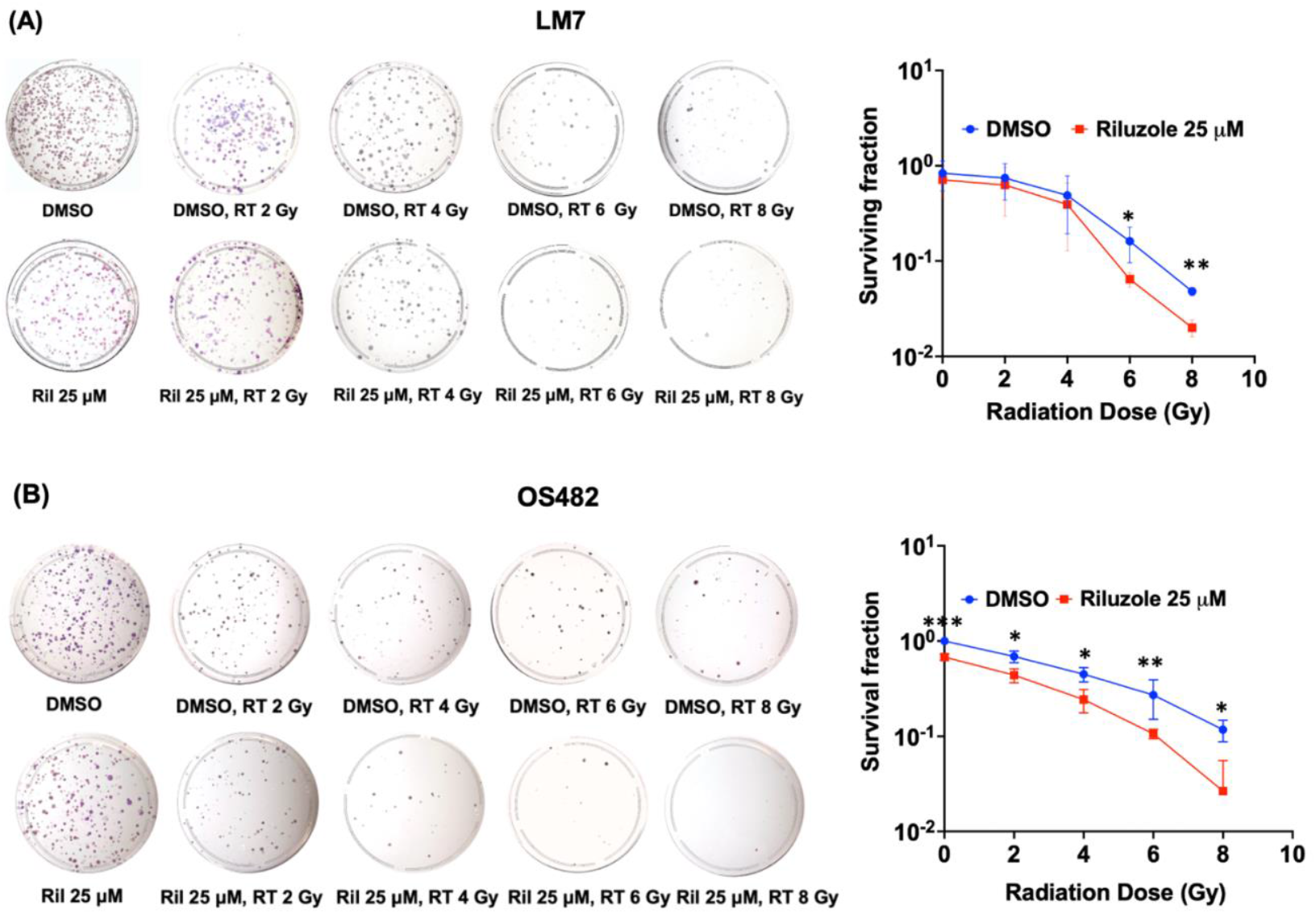
Riluzole pre-treatment decreases survival fraction of irradiated OS cells. **(A)** LM7 and **(B)** OS482 cells were pre-treated with 25 μM Riluzole (Ril) or DMSO (control) for 24 h and were subjected to various dose of X-ray radiation (RT). Ability of Riluzole to affect survival fraction of irradiated cells was assessed using clonogenic assay. The graph shows the dose survival curve for LM7 and OS482 cells treated with Radiation and/or Riluzole. Data are presented as mean ± SD from three independent experiments (*p < 0.05, **p < 0.01).(**p<0.01).

#### Riluzole Elevates SDRT Induced Apoptosis of Osteosarcoma Cells

We next evaluated whether Riluzole could enhance apoptosis in osteosarcoma cells exposed to SDRT. LM7 and OS482 cells were pretreated with 25 μM Riluzole for 24 h prior to SDRT (10 Gy for LM7 ; 6 Gy for OS482). Apoptosis was assessed 24 h post-irradiation using Hoechst 33342 staining. Cells undergoing apoptosis were identified by condensed chromatin and nuclear fragmentation patterns as indicated in red arrows. In LM7 cells, Riluzole alone induced 17.5% apoptosis, while 10 Gy irradiation induced 14.5%. Combination treatment markedly increased apoptosis to 53.5%, demonstrating a synergistic effect **(Fig. 2A)**. In OS482 cells, 6 Gy irradiation induced 25.8% apoptosis, and Riluzole alone induced 34.8%. The combination further enhanced apoptosis to 58%, which was significantly higher than apoptosis induced by each treatment alone **(Fig. 2C)**. These results indicate that Riluzole significantly potentiates radiation-induced apoptosis in osteosarcoma cells, particularly overcoming intrinsic radioresistant phenotype of LM7.

**Figure 2:**
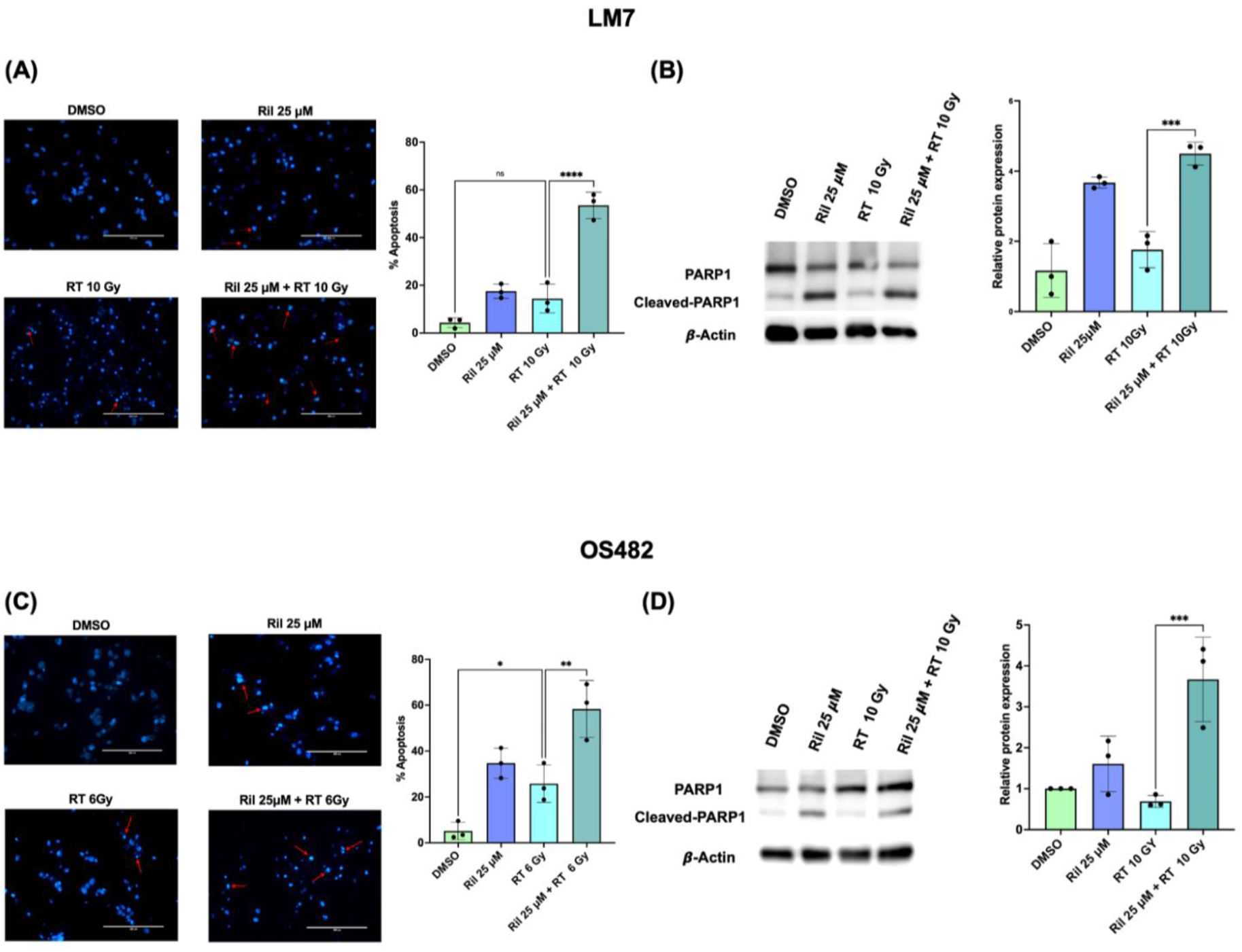
Riluzole promotes SDRT-induced apoptosis in OS cells. **(A)** LM7 and **(C)** OS482 cells were pre-treated with Riluzole (25 μM) or DMSO (control) for 24 h and were subjected to X-ray radiation (10 Gy for LM7; 6 Gy for OS482) or no radiation. Cells were fixed 24 h post irradiation and stained with Hoechst-33258 stain. Apoptotic cells were identified based on nuclear fragmentation and chromatin condensation (indicated by red arrows). Bar graphs show quantification of apoptotic cells from bisbenzimide staining. **(B)** LM7 and **(D)** OS482 cells treated as described above were harvested 5 hours post-irradiation, and western blot analysis was performed to quantify cleaved PARP1 protein levels as a marker of apoptosis. β-actin was used as a loading control. The bar graphs shows relative ratio of cleaved PARP1 to PARP1 protein levels normalized to β-actin. Data are presented as mean ± SD from three independent experiments (*p < 0.05, **p < 0.01, ***p < 0.005).

Cleaved PARP1 is a well-established marker of apoptosis. To further validate these pro-apoptotic effects of the combination treatment, cleaved PARP1 levels were assessed by western blot in LM7 and OS482 cells following SDRT with or without Riluzole pretreatment. Combination treatment significantly increased PARP1 cleavage (p<0.05), compared to irradiation alone at 5 h post-SDRT in both LM7 **(Fig. 2B)** and OS482 cells **(Fig. 2D)** further supporting Riluzole’s pro-apoptotic, radiosensitizing effect.

#### Riluzole Increases Radiation Induced ROS Production

Our previous work has shown that Riluzole induces apoptosis by increasing reactive oxygen species (ROS) in osteosarcoma cells^35^. To assess whether Riluzole potentiates ROS generation following SDRT, intracellular ROS levels were measured post the combination treatment using a fluorescence-based assay. LM7 cells were pretreated with Riluzole (25 μM or 50 μM) or DMSO (control) for 1 h and then irradiated with X-rays (8 Gy for LM7; 6 Gy for OS482). Intracellular ROS levels were quantified 1 h post-irradiation. As shown in **Fig.3A**, SDRT treatment alone does not elicit a significant increase in ROS production in LM7 cells compared to control (DMSO). In contrast, Riluzole (25 μM and 50 μM) pretreatment prior to irradiation resulted in significant elevation of ROS compared to SDRT alone **(3A)**. In OS482 cells, 6 Gy irradiation alone induced a modest increase in ROS, which was further potentiated by Riluzole pretreatment, which can be seen as a synergistic elevation of ROS 1 h post-irradiation (**Fig.3B**). Riluzole induces ROS amplification and augments the oxidative effects of ionizing radiation likely contributing to enhanced DNA damage and radiosensitization.

**Figure 3:**
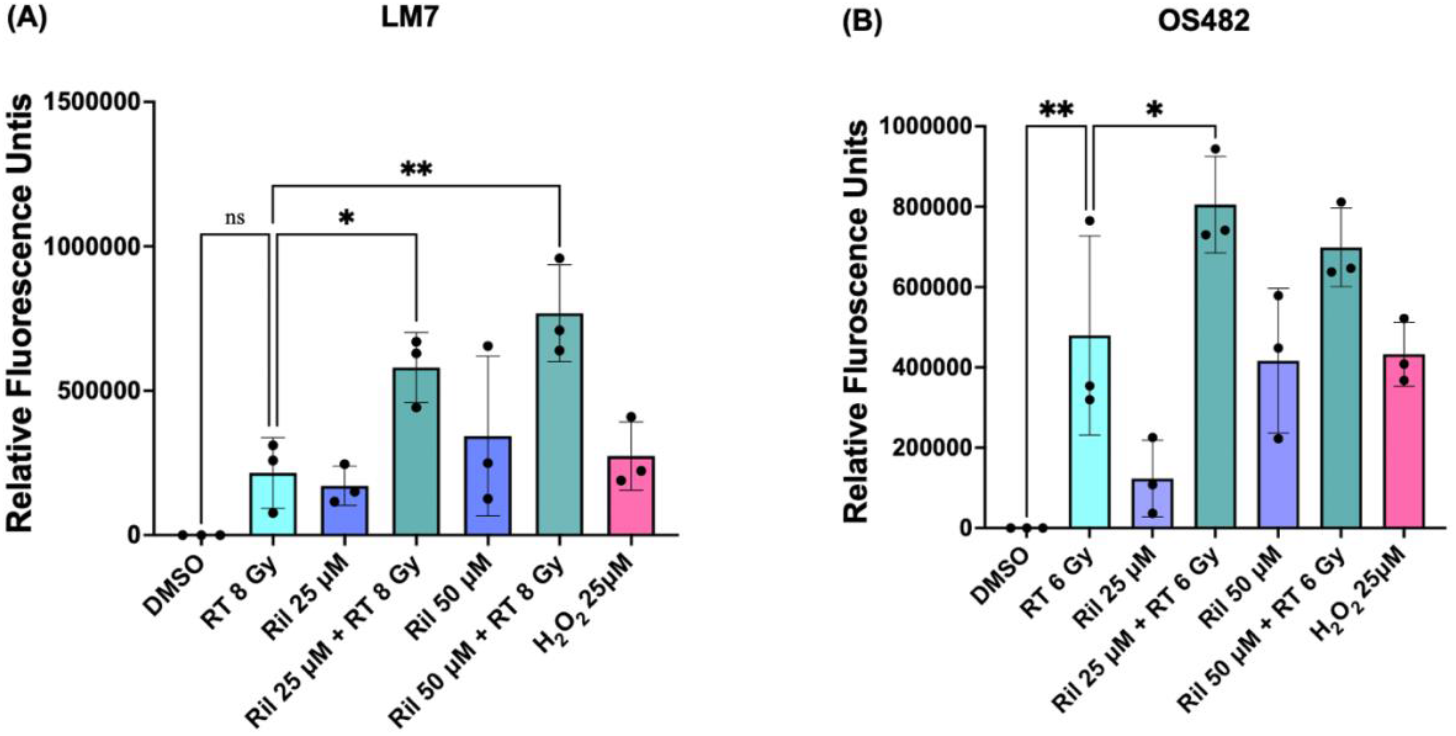
Riluzole promotes radiation-induced ROS production in OS cells. **(A)** LM7 and **(B)** OS482 cells were pre-treated with Riluzole (25 μM, 50 μM)or DMSO for 1 h and then subjected to X-ray radiation (8 Gy for LM7; 6 Gy for OS482) or no radiation. Cells treated with H_2_O_2_ were taken as positive control. ROS levels were measured 1 h post-irradiation using fluorescence based intracellular ROS assay kit. Data are presented as mean ± SD from three independent experiments (*p < 0.05, **p < 0.01).

#### Riluzole Induces G2/M Phase Cell Cycle Arrest

To evaluate the effect of Riluzole on cell cycle progression, LM7 and OS482 cells were treated with 25 μM Riluzole or DMSO (control) for 24 h and cell cycle analysis was carried out using flow cytometry. Riluzole treatment induced a pronounced G2/M phase arrest (∼3.5 fold) in osteosarcoma cells, with 54.1% of LM7 cells and 47.4% of OS482 cells in G2/M, compared with 14.2% and 14.3% in DMSO-treated controls, respectively (**Fig. 4**). Since cells in G2/M are more sensitive to radiation, Riluzole-induced accumulation of osteosarcoma cells in this phase contributes to its radiosensitizing effect.

**Figure 4:**
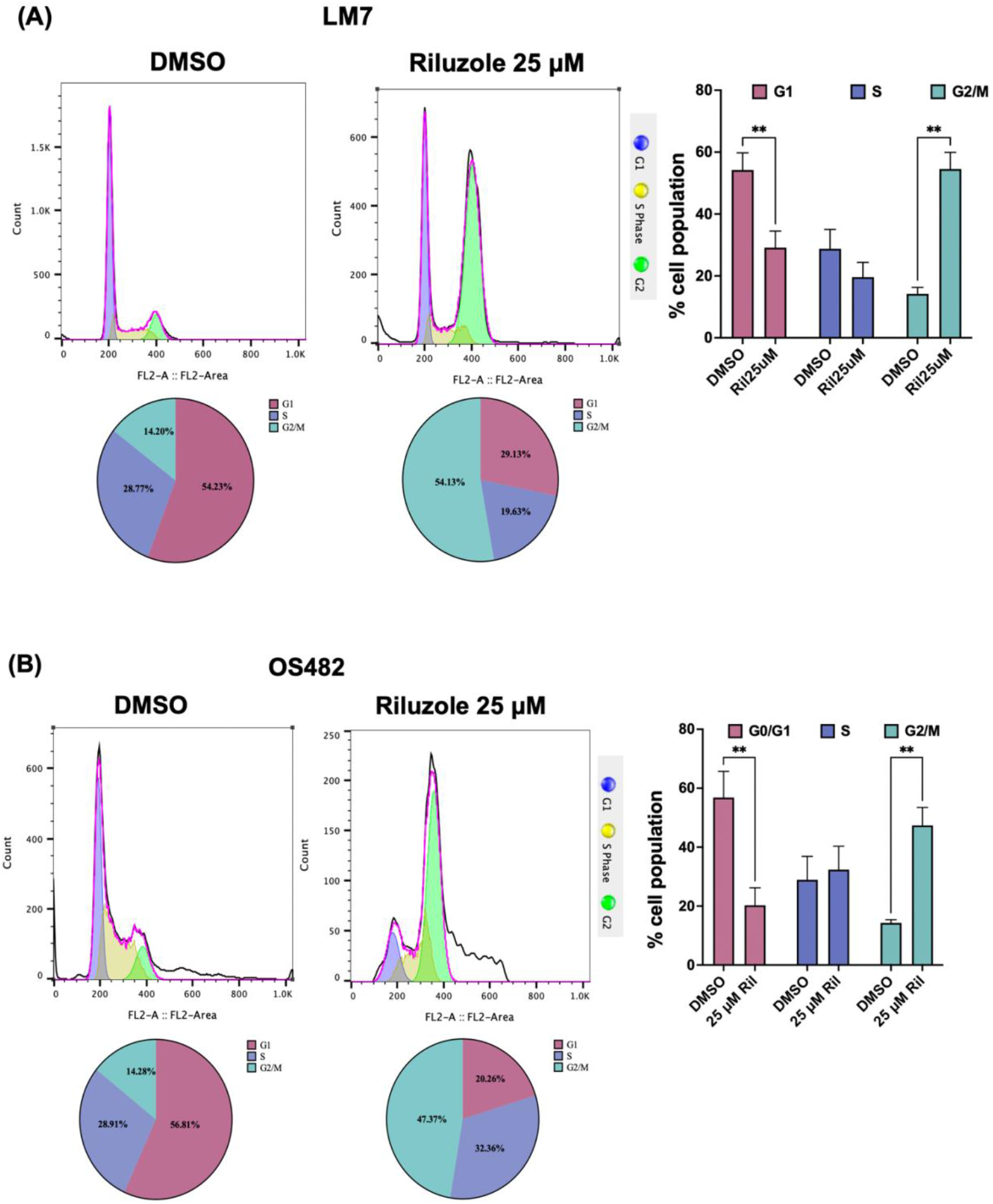
Riluzole induces G2/M phase cell cycle arrest. **(A)** LM7 and **(B)** OS482 cells were pretreated with Riluzole (25 μM) or DMSO (control) for 24 h. Cells were harvested 24 h after irradiation, fixed, and stained with propidium iodide (PI) for DNA content analysis. Cell cycle distribution was evaluated by flow cytometry. The bar graphs show quantification of cells in various phases of cell cycle in LM7 and OS482 cells based on flow cytometric analysis. Data are presented as mean ± SD from three independent experiments (*p<0.05, **p<0.01).

#### Riluzole Potentiates Radiation-Induced DNA Damage

Double-stranded DNA breaks induced by irradiation are marked by accumulation of γ-H2AX foci. To evaluate whether Riluzole enhances SDRT-induced DNA damage, LM7 and OS482 cells were pretreated with Riluzole (25 μM) and subjected to SDRT (10 Gy for LM7; 6 Gy for OS482). γ-H2AX protein levels were quantified by western blotting. Riluzole pretreatment significantly increased γ-H2AX accumulation compared with either treatment alone in both LM7 **(Fig. 5A)** and OS482 **(Fig. 5B)** cells. These results indicates that Riluzole enhances radiation-induced DNA damage in osteosarcoma cells.

**Figure 5:**
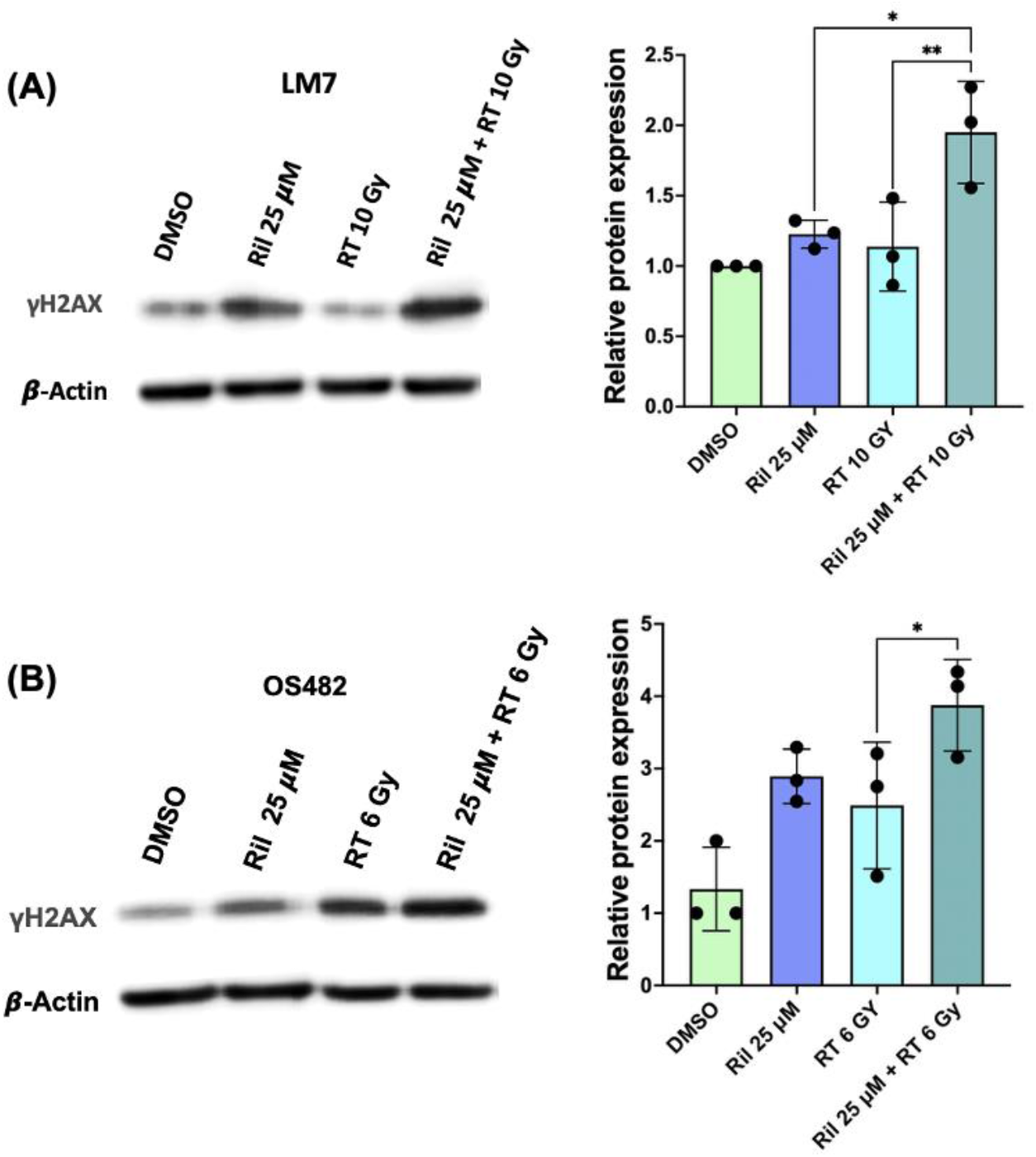
Riluzole enhances γ-H2AX accumulation in irradiated OS cells. **(A)** LM7 and **(B)** OS482 cells were pretreated with Riluzole (25 μM) or DMSO (control) for 24 hours, followed by exposure to X-ray radiation (10 Gy for LM7; 6 Gy for OS482) or no radiation. Cells were harvested 1-h post-irradiation, and γ-H2AX levels were analyzed by Western blotting as a marker of DNA double-strand break (DSB) formation. β-actin served as a loading control. The bar graph shows relative γ-H2AX protein levels normalized to β-actin, quantified from Western blot analysis. Data are presented as mean ± SD from three independent experiments (*p<0.05, **p<0.01).

#### Riluzole Inhibits Radiation-Induced PARP1 Activity

PARP1 is a key regulator of DNA damage repair, and its activation is reflected by increased PARylation, which facilitates recruitment of DNA repair proteins to damaged sites. To assess whether Riluzole modulates PARP1 activity, PARylation levels were measured following SDRT with or without Riluzole pretreatment in LM7 cells. SDRT alone induced increase in PRAP1 PARylation, indicating activation of PARP1 and subsequent activation of DNA repair pathways. Importantly, Riluzole effectively suppressed the radiation-induced PARP1 activation, as evidenced by marked reduction in PARylated PARP1 levels in the combination group **(Fig. 6)**. These findings suggest that Riluzole impairs radiation induced DNA damage repair, as consistent with the elevated γ-H2AX levels observed, thereby reinforcing its potential as a radiosensitizer.

**Figure 6:**
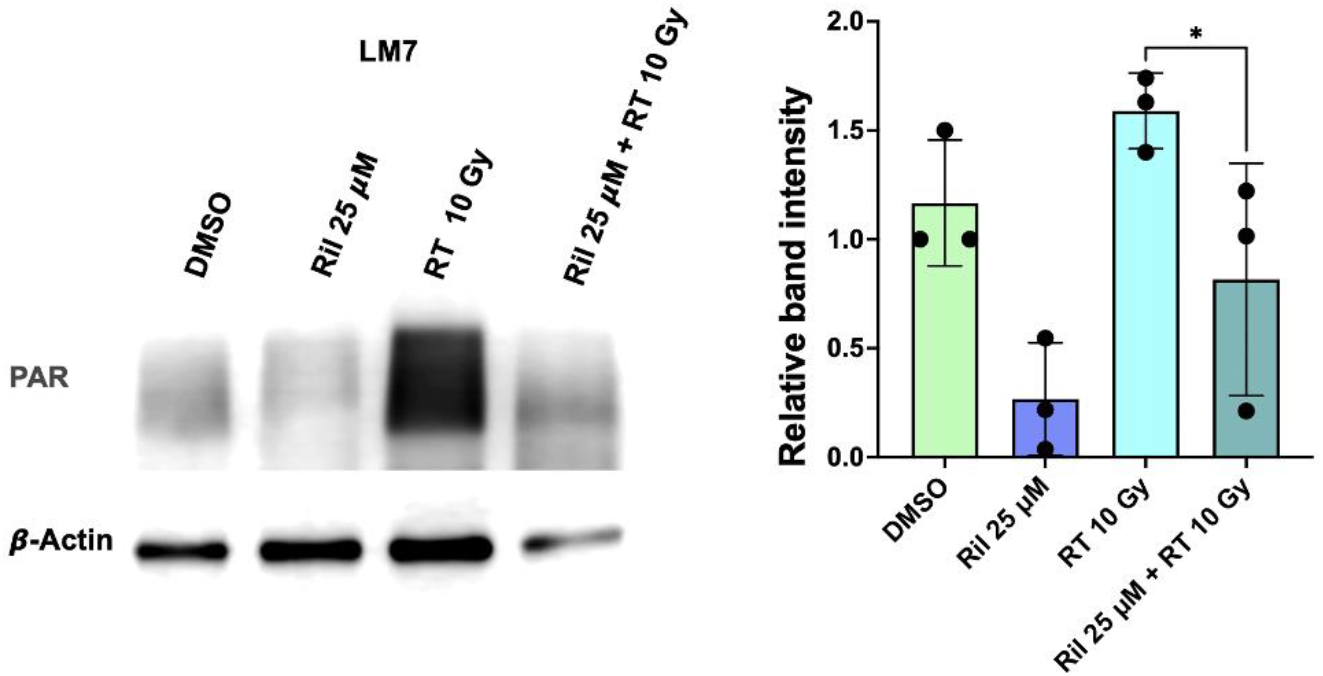
Riluzole decreases radiation-induced PARylated PARP1 levels in LM7 cells. LM7 cells were pretreated with Riluzole (25 μM) or DMSO for 24 h and exposed to X-ray radiation 10 Gy or no radiation. Cells were harvested 24 h post-irradiation, and PARylated PARP1 levels were analyzed by Western blotting. β-actin was used as a loading control. The bar graph shows relative expression of PARylated PARP1 normalized to β-actin, quantified from western blot analysis. Data are presented as mean ± SD from three independent experiments (*p<0.05).

### B. Riluzole Suppresses VEGFA Expression and Modulates Pro-Angiogenic Signaling in Osteosarcoma Cells

#### Riluzole Suppresses Basal and Radiation-stimulated VEGFA Expression

Osteosarcoma cell lines demonstrate robust VEGFA protein expression compared to human fetal osteoblast (hFOB) cells, highlighting the pro-angiogenic phenotype of these tumor cells **(Fig. 7A)**. Treatment with Riluzole significantly reduced basal VEGFA levels compared to DMSO-treated controls, suggesting its potential to impair tumor angiogenesis **(Fig. 7B)**. Furthermore, exposure to SDRT upregulated VEGFA gene expression in osteosarcoma cells, consistent with a compensatory pro-survival angiogenic response to DNA damage **(Fig. 7C)**. Notably, Riluzole effectively suppressed this radiation-induced VEGFA protein upregulation in both LM7 **(Fig. 7D)** and OS482 cells **(Fig. 7E)**. This finding indicates the potential of Riluzole to counteract angiogenic signaling that contributes to radiation resistance.

**Figure 7:**
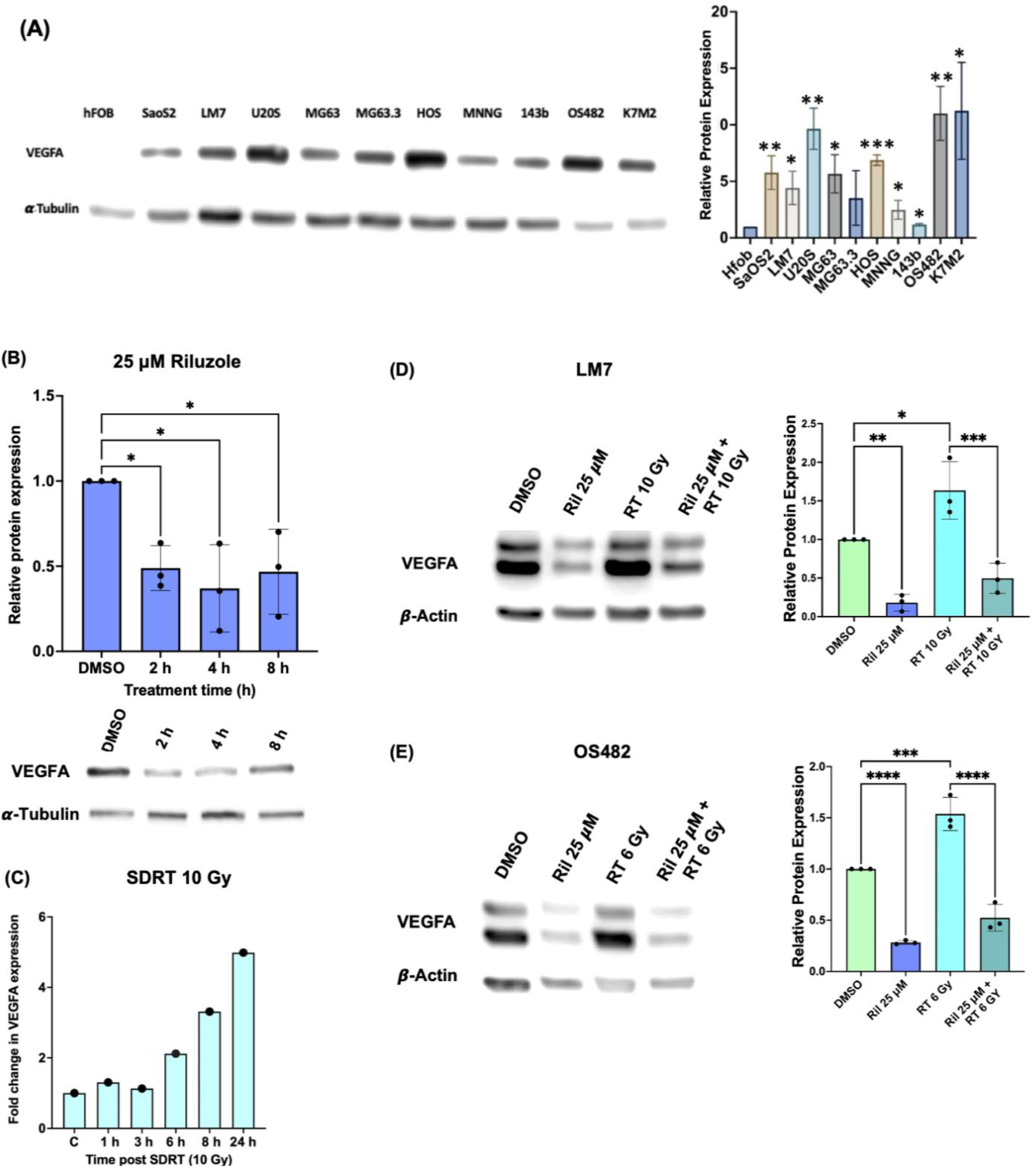
Riluzole suppresses VEGFA expression in osteosarcoma (OS) cells. **(A)** VEGFA protein levels in osteosarcoma cell lines relative to human fetal osteoblast cells (hFOB), assessed by Western blotting. β-actin was used as a loading control. The Bar graph show VEGFA levels normalized to β-actin, quantified from western blot analysis. **(B)** LM7 cells were pretreated with Riluzole (25 μM or 50 μM) or DMSO (control) for the indicated times, and VEGFA protein expression was analyzed by Western blotting. α-Tubulin was used as a loading control. The bar graph shoes VEGFA protein expression normalized to α-Tubulin, quantified from western blot analysis **(C)** LM7 cells were exposed to SDRT (10 Gy), and relative VEGFA mRNA expression was measured by RT-qPCR at the indicated times post-irradiation. **(D)** LM7 and **(E)** OS482 cells were pretreated with Riluzole (25 μM) or DMSO (control) for 24 h, followed by X-ray irradiation (10 Gy for LM7; 6 Gy for OS482) or no irradiation. VEGFA protein levels were determined by Western blotting 24 h post-irradiation. β-actin was used as a loading control. The bar graphs show VEGFA levels normalized to β-actin. Data represent mean ± SD from three independent experiments (*p<0.05, **p<0.01, ***p<0.001, ****p<0.0001).

## Materials and Methods

Riluzole (Cat. #0768) was purchased from Tocris Bioscience. Bisbenzamide (Hoechst 333258; Cat. #382065) was obtained from Millipore Sigma. Protease–phosphatase inhibitor cocktail (Cat. #78442), enhanced chemiluminescence (ECL) reagents (Cat. #34577), propidium iodide (Cat. #P3566), crystal violet (Cat. #C581 - 25), and paraformaldehyde, 32% w/v aqueous solution (Cat. #473779M) and SYBR® Green (Cat. #A25741) were purchased from Thermo Fisher Scientific, USA. RNeasy® mini kit (Cat. #74104) was purchased from Qiagen, Germany. The OxiSelect™ Intracellular ROS Assay Kit was purchased from Cell Biolabs, Inc. Irradiation was performed using a Philips MG-324 X-ray irradiator, delivering the desired dose at 117 cGy/min. Flow cytometry analysis was carried using FACSCalibur (Becton-Dickinson Biosciences). Western blots were analyzed using a LI-COR C-DiGit® Blot Scanner.

### Antibodies and Primers

The following primary antibodies were used for Western blotting: anti-Poly/Mono-ADP Ribose (Cell Signaling, Cat. #83732), anti-phospho-H2A.X (Ser139) (Cell Signaling Technology, Cat. #2577), anti-β-actin (Cell Signaling Technology, Cat. #3700), anti-α-Tubulin (Cell Signaling Technology, Cat. #3873), and anti-VEGFA (Abcam, Cat. #ab46154). HRP-conjugated secondary antibodies were purchased from Cell Signaling Technology (anti-rabbit-HRP, Cat. #7074; anti-mouse-HRP, Cat. #7076S). Customized human VEGFA and GAPDH primers were obtained from Sigma.

Human VEGFA: Forward 5’ ATGATTCTGCCCTCCTCCTT 3’

Reverse: 5’ATGATTCTGCCCTCCTCCTT 3’

Human GAPDH: Forward: 5’ GGTCTCCTCTGACTTCAACA 3’

Reverse: 5’ AGCCAAATTCGTTGTCATA 3’

### Cell culture

LM7 (human osteosarcoma)^40^ and OS482 (mouse osteosarcoma) cells^41^ were cultured in DMEM supplemented with 10% fetal bovine serum (FBS), 1 mM pyruvate, 2 mmol/l GlutaMAX-I, 100 U/mL penicillin, 100 µg/mL streptomycin, in a humidified incubator maintained at 37 °C with 5% CO_2_.

### Bis-benzamide (Hoechst 333258) staining

Fluorochrome bis-benzamide (Hoechst 333258) staining was used to visualize morphological changes associated with apoptosis. LM7 and OS482 (1 × 10^6^) cells were treated with 25 µM Riluzole for 24 h and subjected to single-dose irradiation (10 Gy for LM7 and 6 Gy for OS482). At 24 h post-irradiation, cells were fixed with 4% paraformaldehyde and stained with 24 µg/mL bis-benzamide trihydrochloride for 10 minutes. Apoptotic cells were visualized and quantified using a fluorescence microscope equipped with a DAPI filter. A minimum of 500 cells were analyzed per treatment group.

### Clonogenic assay

Ability of Riluzole to effect survival fraction of irradiated osteosarcoma cells was evaluated using Clonogenic assay. LM7 and OS482 cells (1 × 10^6^) were treated with 25 µM Riluzole for 24 h and irradiated with X-rays (2 Gy, 4 Gy, 6 Gy, or 8 Gy). Following treatment, cells were trypsinized and seeded at a density of 1000 cells per 100*10mm culture dish and cultured for 10 days to allow colony formation. The colonies were fixed with 4% PFA and stained using 0.05% crystal violet for 5 mins and counted manually. Colonies containing ≥50 cells were scored as survivors. Plating efficiency (PE) was calculated from control (DMSO) plate, and survival fractions (SF) were determined relative to this efficiency.

% PE = (No. of Colonies in control plate/ No. of cells seeded)*100

SF= (No. of Colonies / No. of cells seeded * PE)

### Intracellular Reactive oxygen species (ROS) assay

The ability of Riluzole to enhance SDRT-induced oxidative stress was assessed using an intracellular ROS detection assay kit. LM7 cells were seeded in 24-well plates (30,000 cells/well), treated with Riluzole (25 or 50 µM) or DMSO (control) for 1 h, and irradiated with 8 Gy (LM7) or 6 Gy (OS482). Cells treated with 25 μM H_2_O_2_ served as positive controls. Cells were harvested after for 1 h of incubation at 37 °C and Intracellular ROS levels were quantified using the OxiSelect Intracellular ROS Assay Kit according to the manufacturer’s instructions. Briefly, cells were washed with PBS and incubated for 1 h with the fluorogenic probe 2′,7′-dichlorodihydrofluorescein diacetate (DCFH-DA) in serum-free DMEM media. The cells were treated with 100 μM H2O2 for 1 hour as a positive control. Following incubation, cells were washed and lysed in 1× lysis buffer and fluorescence intensity was measured at 480/530 nm using a SpectraMax i3 plate reader.

### Cell cycle analysis

The effect of Riluzole on cell cycle distribution was assessed using flow cytometry. LM7 and OS482 (1 × 10^6^) cells were seeded in 100mm tissue culture plate and serum-starved for 16 h, after which the medium was replaced with complete growth medium containing 25 µM Riluzole or DMSO (control) for 24 h. Following treatment, cells were harvested by trypsinization, washed twice with PBS, treated with 50 ul of RNAse A solution (1mg/mL; Sigma, R-4875) and fixed in ethanol overnight. The cells were then stained with propidium iodide (PI) (1mg/mL; Sigma P4170). DNA content was then quantified by flow cytometry on FACSCantoII to determine the proportion of cells in different cell cycle phases.

### Quantitative Reverse Transcription-Polymerase Chain Reaction (RT-qPCR)

Effect of SDRT on VEGFA gene expression in osteosarcoma cells was as measured using RT-qPCR. LM7 cells (1*10^6^ cells) were irradiated with x-ray (10 Gy; delivered at 117 cGy/min). Following the treatment, total RNA from the cells were extracted using RNeasy® mini kit following the manufacturers instruction and eluted in 30uL RNAase free water. QuantiTect® reverse transcription kit was then used to convert 1 ug of RNA to cDNA. To quantify the VEGFA mRNA levels in the cells, the cDNA template was mixed with SYBR® Green and primer mix consisting of 0.4 μM Forward primer, 0.4 μM Reverse primer. The cycle setting was 95 °C for 10 mins (15 s at 95 °C for denaturation, 1 min. at 60 °C for annealing, and 1 min. at 72 °C for extension) for 40 cycles. The cycle threshold (C_t_) values of VEGFA were normalized to C_t_ of GAPDH for LM7 and OS482 cells to obtain the relative threshold values and the fold change in VEGFA expression was calculated.

### Western blot analysis

Cells treated with Riluzole and/or irradiation were lysed on ice using RIPA buffer (50mM Tris, 150mM NaCl, 1mM EDTA, pH 8.0, 1% NP40, 5% glycerol, 1mM Dithiothreitol) supplemented with protease–phosphatase inhibitor cocktail. Lysates were sonicated (20% amplitude, 10 s) and centrifuged at 13,500 × g for 30 min at 4 °C. Protein concentrations were determined using a bicinchoninic acid (BCA) assay kit. Equal amounts of protein were resolved by SDS–PAGE and transferred onto PVDF membranes. Membranes were blocked using 5% non-fat dry milk and incubated overnight at 4 °C with primary antibodies against proteins of interest, followed by HRP-conjugated secondary antibodies. β-actin or α-tubulin was used as loading control. Protein bands were detected using enhanced chemiluminescence and imaged with LiCOR digital system.

### Statical analysis

All the experiments were performed in triplicates. Results from RT-qPCR is from single experiment. Statistical analyses was performed using GraphPad Prism 10.0. Data are presented as mean ± standard deviation. Differences between groups were evaluated using either one-way ANOVA or unpaired Student’s two-tailed t-test. p-value < 0.05 was considered statistically significant.

## Discussion

Our study establishes Riluzole as a potent radiosensitizer for osteosarcoma, capable of enhancing the efficacy of SDRT. Riluzole overcomes two major barriers to effective radiation therapy in osteosarcoma: (1) it enhances tumor cell radiosensitivity, thereby potentially reducing the dose of single-dose radiation therapy required to achieve therapeutic efficacy, and (2) it suppresses the expression pro-survival angiogenic factor VEGFA, which may indirectly diminish endothelial survival signaling. These findings highlight the potential of Riluzole to improve radiation outcomes in osteosarcoma and provide a strong rationale for its clinical translation as a radiosensitizing agent.

Osteosarcoma is an extremely radioresistant tumor, restricting the clinical efficacy of radiotherapy despite advances in delivery techniques^5,30,40^. While SDRT has demonstrated promise in hypervascular tumors^11,29^, its effectiveness in osteosarcoma is constrained by toxicity to adjacent critical structures and radiation-induced pro-survival signaling in the tumor microenvironment^28,29,42^. Previous strategies that selectively targeted either tumor cells or vasculature achieved only modest success, highlight the need for dual-targeted radiosensitizers, such as Riluzole, that can overcome this bidirectional crosstalk between compartments.

At the cellular level, Riluzole significantly enhanced SDRT efficacy, reducing clonogenic survival in LM7 and OS482 cells. The combination synergistically increased apoptosis in LM7 and additively in OS482 cells, as confirmed by Hoechst-333258 staining and cleaved PARP1 levels, suggesting that Riluzole effectively radiosensitizes osteosarcoma cells *in vitro*.

Approximately two-thirds of radiation-induced DNA damage is mediated by reactive oxygen species (ROS)^43^. Cancer cells counteract oxidative stress via robust antioxidant defenses such as glutathione (GSH)^44,45^. Radiosensitizers that impair this antioxidant systems improve oxidative damage and thereby enhance the efficacy of ionizing radiation. Riluzole has been shown to inhibit xCT (SLC7A11), the cystine/glutamate antiporter, depleting intracellular GSH and impairing redox homeostasis^15^. In this study, Riluzole amplified ROS generation in osteosarcoma cells following SDRT, consistent with a pro-oxidant radiosensitizing effect. While SDRT alone induced minimal ROS in LM7 cells, Riluzole pretreatment markedly elevated ROS induced by SDRT. Similarly, in OS482 cells, Riluzole combined with SDRT synergistically increased ROS production, further supporting its role in promoting oxidative stress in irradiated cells.

Reactive oxygen species (ROS) accumulation is closely linked to the induction of DNA double-strand breaks (DSBs), one of the most lethal forms of DNA damage if left unrepaired. Previous studies have demonstrated that Riluzole enhances radiation-induced DNA damage in glioma and melanoma cells^34^. In osteosarcoma cells, we observed that Riluzole induces pronounced G2/M cell cycle arrest, a phase in which cells are inherently more radiosensitive due to reduced DNA repair efficiency^46^. Consistent with this, γ-H2AX, a well-established marker of DSBs, was significantly elevated in irradiated osteosarcoma cells pretreated with Riluzole compared to either treatment alone, indicating that Riluzole amplifies SDRT-induced DNA damage. Mechanistically, this radiosensitization is likely mediated through a combination of enhanced ROS production, impaired DNA repair processes, and cell cycle redistribution, collectively leading to increased apoptosis and improved therapeutic efficacy of SDRT.

Another mechanism of radioresistance in osteosarcoma is the extraordinary DNA repair capability of these cells, particularly through PARP1 which resolves single-strand breaks and coordinates recruitment of DNA repair machinery^47^. PARP1 inhibitors have been explored as radiosensitizers in multiple tumor types, highlighting the therapeutic potential of targeting DNA repair pathways^9,48^. Here, we demonstrate for the first time that Riluzole suppressed PARP1 activity in osteosarcoma cells, as evidenced by reduced PARylation of PARP1 in Riluzole treated cells, thereby impairing DNA repair and allowing accumulation of unrepaired DSBs. When combined with Riluzole-induced ROS generation and G2/M cell cycle arrest, PARP1 inhibition synergistically amplifies DNA damage, leading to enhanced apoptosis and radiosensitization. These findings establish a mechanism through which Riluzole overcomes intrinsic tumor radioresistance, supporting its potential as a potent radiosensitizer for osteosarcoma.

Beyond tumor-intrinsic mechanisms, Riluzole also modulates the angiogenic compartment. VEGFA, a critical regulator of angiogenesis, is among the most highly expressed pro-angiogenic factor in osteosarcoma^23,49^. Following SDRT, VEGFA expression is further upregulated in tumor cells, promoting endothelial survival by inhibiting acid sphingomyelinase (ASMase)–mediated, ceramide-driven apoptosis through activation of the VEGFA/VEGF Receptor signaling axis ^21,29,42^. Consistent with this, we observed that SDRT increased VEGFA expression in osteosarcoma cells, potentially reinforcing tumor–endothelial crosstalk and compensatory pro-survival signaling within the tumor microenvironment. Notably, Riluzole suppressed both basal and SDRT-induced VEGFA expression, suggesting that it may indirectly attenuate VEGFA/VEGF receptor-mediated endothelial survival, thus promote optimal response to SDRT in these cells. This is particularly significant in the context of SDRT, whose therapeutic efficacy is often hindered by VEGFA-driven angiogenic radioresistance ^11,29^. This dual targeting of tumor and vascular compartments uniquely positions Riluzole as a potent radiosensitizer capable of overcoming complimentary mechanisms of radioresistance.

Collectively, these results establish Riluzole as an effective radiosensitizer for osteosarcoma in combination with SDRT. Through enhanced ROS generation, G2/M arrest, amplified DNA damage, inhibition of PARP1-mediated repair, and suppression of VEGFA expression, Riluzole simultaneously targets tumor-intrinsic and microvasculature-mediated resistance. Given its favorable safety profile and existing FDA approval, Riluzole represents a clinically translatable strategy to improve the therapeutic index of SDRT in osteosarcoma.

## Acknowledgements

We thank Eugenie E. Kleinerman from the MD Anderson Cancer Center, Houston, Texas, for the generous gift of LM7 cells.

